# Consequences of recombination for the evolution of the mating type locus in *Chlamydomonas reinhardtii*

**DOI:** 10.1101/565275

**Authors:** Ahmed R. Hasan, Jaspreet K. Duggal, Rob W. Ness

**Author notes:** Correspondence: Ahmed Hasan.

## Abstract

**Rationale:** Recombination suppression in sex chromosomes and mating type loci can lead to degeneration due to reduced selection efficacy and Muller’s ratchet effects. However, genetic exchange in the form of non-crossover gene conversions may still take place within crossover-suppressed regions. Recent work has found evidence that gene conversion may explain the low levels of allelic differentiation in the dimorphic mating type locus (*MT*) of the isogamous alga *Chlamydomonas reinhardtii*. However, no one has tested whether gene conversion is sufficient to avoid the degeneration of functional sequence within *MT*.

**Methods:** Here, we calculate levels of linkage disequilibrium (LD) across *MT* as a proxy for recombination rate and investigate its relationship to patterns of population genetic variation and the efficacy of selection in the region.

**Results:** We find that levels of LD predict selection efficacy across *MT*, and that purifying selection is stronger in shared genes than *MT*-limited genes to the point of being equivalent to that of autosomal genes.

**Conclusions:** We argue that isogamous systems without secondary sexual characteristics exhibit reduced selective pressure to differentiate sex chromosomes, and that recombination via gene conversion plays an important role in both reducing differentiation and preventing degeneration of crossover suppressed mating type loci.

## Introduction

Sexual reproduction is widespread across both unicellular and multicellular eukaryotes primarily to shuffle genetic material between compatible mates. Despite the role of sex in promoting recombination and therefore improving the efficacy of selection in the genome, evidence and theory suggest that sex or mating-type determining regions themselves are often recombination-suppressed (Bachtrog et al., 2011, Abbott et al., 2017). Recombination suppression in these regions is thought to be a result of selection to preserve linkage between sex-determining loci that and sexually antagonistic alleles (Charlesworth, 1996; but see Branco et al. 2017 and Ponnikas et al. 2018). Through time, the region over which recombination is suppressed can expand and potentially form complete sex chromosomes. The suppression of recombination in mating-type determining regions or chromosomes has major consequences for their molecular evolution relative to the remainder of the autosomal genome and poses questions about how their function is maintained over evolutionary time (Bergero and Charlesworth, 2009, Wright et al., 2016).

Y-chromosome-degeneration is one of most well-known and best-documented cases of the altered evolutionary trajectory of sex chromosomes. Y-chromosomes have undergone substantial erosion of gene content and has accumulated repetitive elements in many taxa (Bachtrog, 2006, 2013), such as plants (Ming et al., 2011), mammals (Graves, 2006), birds (Berlin and Ellegren, 2006), and insects (Singh et al., 2014). Y-chromosome degeneration is believed to be a consequence of the suppression of recombination that defines these sex determining regions. One major consequence of reduced recombination is the Hill-Robertson effect, wherein selection is less effective due to interference between selected loci at linked sites (Hill and Robertson, 1966, Charlesworth and Charlesworth, 2000, Hough et al., 2017). Additionally, Muller’s ratchet results in the irreversible accumulation of deleterious mutations due to stochastic loss of the least mutated haplotype (Muller, 1964, Felsenstein, 1974, Gordo and Charlesworth, 2001). Lastly, recombination suppression can result in reduced efficacy of selection against structural mutations, including transposable element insertions, gene loss, and chromosomal inversions. Such structural mutations, particularly chromosomal inversions, also drive further reductions in recombination by disrupting pairing of homologous chromosomes in meiosis and can therefore expand the boundaries of sex-or mating type-determining regions (Lahn and Page, 1999, Kirkpatrick, 2010, Wright et al., 2016).

The distinction between the sexes ultimately stems from the difference in the size and investment into their respective gametes (Kodric-Brown and Brown, 1987). In such anisogamous systems, selective pressure for secondary sexual characteristics has the potential to favour sexually antagonistic alleles and drive the evolution of completely differentiated sex chromosomes (Bell, 1978, Charlesworth, 1978). In isogamous systems (i.e. with equal sized gametes), compatible mates are known as mating-types (Charlesworth, 1978, Hoekstra, 1987); the genetic loci that determine these mating-types are often smaller than complete chromosomes, but still frequently involve the suppression of recombination and subsequent genetic differentiation (Fraser and Heitman, 2004). Many well-documented examples of mating type locus evolution come from Volvocine green algae (Chlorophyta) which include the isogamous unicellular model organism *Chlamydomonas reinhardtii* and the multicellular anisogamous alga *Volvox carteri* (Umen, 2011, Umen and Olson, 2012). Unlike mating types from familiar yeast species (Dujon, 2010), mating type can not switch in *C. reinhardtii*, and is inherited as a single Mendelian trait (Goodenough et al., 2007). In *C. reinhardtii*, mating types are determined by a haploid, genetically encoded *∼* 500 kb mating type locus (henceforth *MT*) on chromosome 6, whose allelic state encodes an individual as either *MT+* and *MT–*. *MT* is subdivided into three domains: the centremost is known as the R domain, where recombination is suppressed. The R domain carries chromosomal inversions and repetitive regions (Ferris and Goodenough, 1994) as well as numerous genes that are common to both *MT* alleles. There are also a small number of *MT*-limited genes, including the *MT–* gene *mid* that determines mating type specificity and the *MT+* gene *fus1* that facilitates fusion of compatible gametes (Ferris and Goodenough, 1997, Lin and Goodenough, 2007). Flanking the R domain are the telomere-proximal and centromere-proximal domains (T and C domains, respectively), which unlike the R domain are relatively syntenic (Ferris et al., 2002, De Hoff et al., 2013). Interestingly, *MT* also ensures organelle inheritance in *C. reinhardtii* : following sex between two haploid gametes, a diploid zygote forms, in which degradation of the chloroplast genome inherited from the *MT–* parent and the mitochondrial genome from the *MT+* parent occurs prior to the final meiotic event, thus causing uniparental inheritance (Nishimura et al., 1999, Goodenough et al., 2007, Nishimura, 2010). However, both experimental and population genetic evidence suggests that the chloroplast genome may undergo recombination during meiosis and that uniparental inheritance may be leaky (Sager, 1954, Gillham, 1969, Ness et al., 2016).

Although crossing over within the *MT* locus is suppressed, the three domains (T, R, and C) altogether contain 57 shared homologous genes. In a recent genetic analysis of five shared genes, two of them, *PDK1* and *PR46*, were shown to have undergone ‘cryptic recombination’ in the form of gene conversion between *MT* alleles (De Hoff et al., 2013). In this investigation of *MT*, De Hoff and colleagues showed that diversity and divergence between these five shared genes were comparable to those of autosomal genes, suggesting genetic exchange was frequent enough in *MT* to prevent divergence between homologs. From a population genetic perspective, the presence of recombination predicts that selection efficacy may be higher in shared regions despite suppressed crossing-over (Felsenstein, 1974, Comeron et al., 2008) and therefore may provide a mechanism by which *MT* regions can avoid the degeneration observed in many Y-chromosomes. Moreover, we may also expect consequences of gene conversion on base composition through either more efficient selection on codon usage (Hey and Kliman, 2002) or via the process of GC-biased gene conversion (Galtier and Duret, 2001, Marais, 2003). However, to date, no full-scale investigation of the extent of recombination across the *MT* locus has been conducted, nor the consequences of varying recombination across the region.

Here, we investigate the occurrence and extent of linkage disequilibrium across the entire *C. reinhardtii* mating type locus and report an analysis of its effects on the population genetics and molecular evolution of the shared and *MT–*limited regions. Specifically, we address the following questions: 1) What are the spatial patterns of LD across *MT* including the flanks, T, R and C domains? 2) Do patterns of polymorphism or divergence between *MT* alleles reflect the strength of LD? 3) Do shared and *MT*-limited regions show evidence for differences in selection efficacy? 4) Do patterns of inter-chromosomal LD between mating type alleles and organelle genomes suggest leaky uniparental inheritance? To answer these questions, we characterize the landscape of linkage disequilibrium (LD) across shared regions and relate it to patterns of polymorphism within and surrounding coding sequence from a population genomic data set.

## Materials and Methods

### Strains, sequencing, and alignment

We used whole genome sequence data from 19 (9 *MT+*, 10 *MT–*) natural strains of *Chlamydomonas reinhardtii*, sampled from Quebec, Canada. Strains CC-2935, CC-2936, CC-2937, and CC-2938 were obtained from Flowers et al. (2015), while the remainder were originally published in Ness et al. (2016) (Table S1). We used the *C. reinhardtii* v5.3 reference genome (Merchant et al., 2007) but because it is derived from an *MT+* individual and does not include organelles, we appended the chloroplast genome, the mitochondrial genome, and the *MT–* locus (GenBank accession GU814015.1) to allow mapping of reads derived from these regions. We then aligned reads for strains from the two mating types as follows: for *MT+* individuals, we masked the *MT–* locus prior to alignment; then, for *MT–* individuals, we masked the location of the T and R domains of the *MT+* allele in the reference genome (chromosome_6:298298-826737) prior to alignment. Masking ensured alignment of reads to the correct mating type allele. The C domain was not masked as it is completely syntenic across both alleles (De Hoff et al., 2013) and is not included in the *MT–* sequence. We used the GATK v3.3 tools HaplotypeCaller and GenotypeGVCFs for variant calling (non default settings: `ploidy=1, includeNonVariantSites=true, heterozygosity=0.02, indel_heterozygosity=0.002`).

### Identification and alignment of shared regions

To identify shared regions between the two mating type loci, we used the pairwise aligner LASTZ 1.04 (Harris, 2007) on FASTA files containing the reference sequences of T and R domains across both mating type alleles. The *MT+* locus was assigned as the target and the *MT–* the query. We used a score threshold of 30000; default parameters were otherwise retained. Cases of multiple *MT–* hits to the same *MT+* region, defined as two or more alignments with at least 75% overlap in their *MT+* sequence coverage, were resolved by selecting the highest-scoring match. All alignments were visually inspected using the LASTZ visualizer plugin in Geneious 11.1.4 (www.geneious.org). Using the *MT+* allele (which is approximately 200 kb longer than the *MT–* allele) positions as a reference, we concatenated individually aligned segments into a single FASTA alignment the length of the *MT+* allele that contained only shared regions across all *MT+* and *MT–* strains as output by LASTZ. In this merged alignment, non-shared regions were instead denoted as N across both alleles to preserve the chromosomal positions of shared regions relative to the *MT+* reference. Finally, because the C domain of the *C. reinhardtii* mating type locus is absent from the *MT–* sequence and syntenic across both mating type alleles (Ferris and Goodenough, 1997, De Hoff et al., 2013) we directly appended it to the 3’ end of the merged alignment. *Recombination rate estimation.* To estimate recombination rate variation using patterns of linkage disequilibrium (LD), we estimated levels of disequilibrium using a Python script. Over all shared regions, we calculated the *r*^2^ measure of LD (Hill and Robertson, 1968) for all pairwise combinations of diallelic, non-singleton SNPs within 1 kb of one another. We only considered windows with a minimum of 30 polymorphic sites. Next, we averaged these *r*^2^ values with *Z_nS_* (Kelly, 1997) in non-overlapping 1 kb windows. *Z_nS_* is the average pairwise LD over all variant sites in a given region, defined as follows:

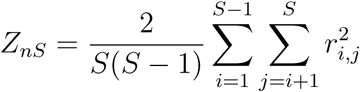

Where *S* is the set of polymorphic sites in the region of interest, *i* and *j* correspond to a pair of variant sites within *S*, and 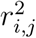 is the square of the correlation coefficient (*r*^2^) between the two loci. Like *r*^2^, *Z*_*nS*_ values range from 0 to 1, with 0 representing linkage equilibrium while a value of 1 indicates complete association. To then visualize broader spatial trends of LD, we then fit a hidden Markov model with three states to *Z*_*nS*_ scores over the mating type locus using the R package depmixS4 (Visser and Speekenbrink, 2010).

### Identification, alignment, and analysis of coding sequence polymorphism

We identified pairs or shared genes from the MT locus alleles using a multi-faceted approach. Firstly, we used the gene IDs present in the genome annotation of *C. reinhardtii* reference v5.3 (Merchant et al., 2007) and the *MT–* sequence. For those genes that were not paired based on gene ID, we performed a reciprocal best BLAST using BLASTn and incorporated shared pairs not already included. Lastly, we resolved a small number of shared pairs using the definitions provided in Table S4 of De Hoff et al. (2013). After excluding the shared genes, we identified *MT*-limited genes as those with no BLAST hit on the opposite *MT* locus. *MT*-limited gene definitions overlapped previous definitions of *MT*-limited genes; however, gene copies from the 16 kb repeat in the *MT+* allele were not included because sequence from these genes was not reliable due to the short-reads used in our study. For comparison to *MT* genes, we also extracted sequence from chromosome 6 genes outside of the *MT* locus and 1000 random genes from the rest of the nuclear genome. After extracting CDS sequences of all individuals from each gene or gene pair, we aligned the sequences based on their translated amino acid sequence using MUSCLE v3.8.31 (Edgar, 2004) and then back translated each to its original nucleotide sequence. All alignments were visually inspected to ensure accuracy. We masked regions where meaningful alignment was impossible, likely due to large insertion/deletion events. Alignments and alignment edits are available at https://github.com/aays/2019-mt-locus.

For each alignment, we calculated population genetic statistics for *MT–* strains, *MT+* strains, and all samples combined. For individual MT groups and *MT*-limited genes, we excluded sites with fewer than 8 called alleles, while for genes shared across both mating type alleles we excluded sites with fewer than 12 alleles called. Within coding sequence, we estimated genetic diversity as *θ_π_* for 0-and 4-fold degenerate sites for each *MT* grouping. We also summarized genetic differentiation within and between *MT* alleles using *F_ST_*, as well as the number of shared, *MT*-private and fixed differences. We also calculated the above population genetic statistics from the same sample groups for genes outside the *MT* locus where recombination is assumed to be present.

### Population genetic consequences of recombination rate variation

We then used our *Z*_*nS*_ scores to investigate the effect of recombination on mating type evolution. First, to examine the effects of recombination on divergence between *MT* alleles, we computed *F*_*ST*_ in non-overlapping 1 kb windows over the shared regions of the *MT* locus, and compared these windowed estimates to the windowed *Z*_*nS*_ values described above.

To then investigate whether recombination affects selection efficacy over the *MT* locus, we computed the ratio of nonsynonymous to synonymous diversity (*π*_*N*_ /*π*_*S*_) and *Z*_*nS*_ over the coding sequences of each shared gene. *π*_*N*_ /*π*_*S*_ values below 1 are indicative of purifying selection, with smaller values suggesting greater selection efficacy.

To test for an effect of recombination on GC-content evolution, either due to selection or GC biased gene conversion, we calculated the GC content of 4-fold degenerate sites (GC4) in *MT*-limited and shared genes. Outside of coding sequence, we calculated the GC content of shared sites annotated as intron or intergenic using the *MT+* reference sequence, to which the alignments were oriented. In *MT*-limited regions, the GC content of intron and intergenic sites was calculated with respect to annotations from the corresponding mating type.

### LD-based analyses of organelle inheritance

To assess leakage in organelle inheritance, we calculated inter-chromosomal LD between each organelle and mating type limited-genes from the corresponding mating type. First, the GATK program HaplotypeCaller was used to generate mating type-separated VCF files containing either of *MT+* individuals and *MT–* individuals. A Python script using PyVCF-0.6.8 was used to compute Lewontin’s D’ statistic (Lewontin, 1964) between organelle genomes and corresponding mating type-specific markers: *mta1* and *fus* in the case of the *MT+*, and *mtd1* and *mid* with the *MT–*. The D’ statistic normalizes values of D between loci, such that a value of 1 always represents complete linkage.

In addition to organelle-to-mating type comparisons, intra-region D’ was calculated in the *MT–* mitochondrial genome with the expectation of complete linkage. Finally, estimates of inter-chromosomal LD across the genome were calculated for use as a null expectation. Given the density of SNPs in the genome of *C. reinhardtii* (*>* 10.2 *×* 10^6^ across the 17 chromosomes), random filtering to 0.02% was applied, so that approximately 1700 arbitrarily selected variants from the entire genome underwent LD calculations for a total of 7.5 *×* 10^5^ pairwise comparisons. Only diallelic and non-singleton SNPs were used in all calculations.

All scripts used in this work can be found at https://github.com/aays/2019-mt-locus. All data analysis and visualization was performed in R (R Core Team, 2018) using the `tidyverse` suite of packages (Wickham, 2017).

## Results

### Breakdown of linkage disequilibrium across the C and T domains

Following alignment of the two *MT* alleles, we calculated LD over the *MT* locus and the remainder of chromosome 6 as a proxy for recombination (i.e. gene conversion), such that regions with higher *Z*_*nS*_ were considered to have lower recombination rates. The distribution of *Z*_*nS*_ values over the *MT* locus and surrounding autosomal sequence is shown in Figure 1. As expected, we observe a strong elevation in LD in the R domain (mean *Z*_*nS*_ = 0.76) as compared to the T domain (mean *Z*_*nS*_ = 0.442) or the C domain (mean *Z*_*nS*_ = 0.473). By contrast, LD levels in the C and T domains approximate autosomal levels (mean chromosome 6 *Z*_*nS*_ = 0.359), suggesting more frequent recombination in these regions.

**Figure 1:**
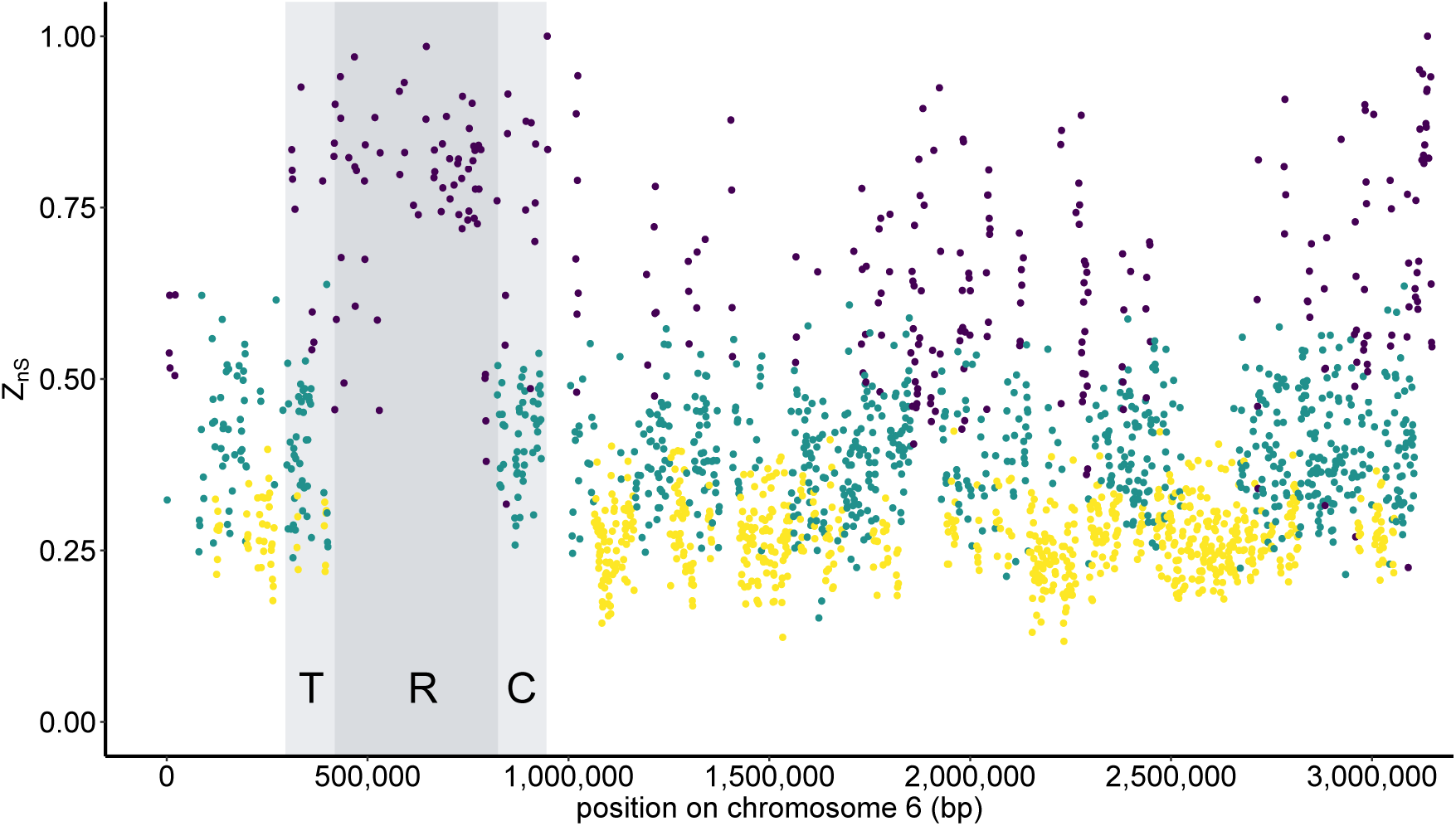
Variation in *Z*_*nS*_ over chromosome 6. Each point represents LD in a 1 kb window of shared sequence across both *MT* alleles. A hidden Markov model with three states was fit to the data to better visualize spatial trends; states are represented with point colouration.

**Figure 2:**
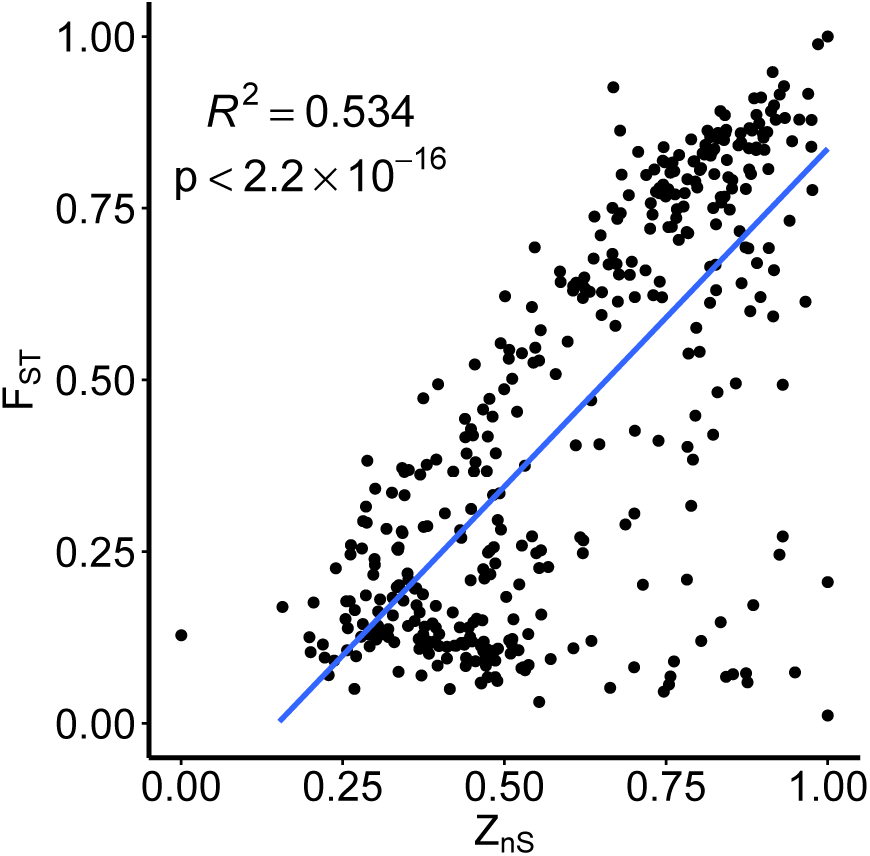
Relationship between differentiation (*F*_*ST*_) and LD (*Z*_*nS*_). Each point represents a 1 kb window of shared sequence between both *MT* alleles.

### Differentiation between shared regions inversely scales with recombination rate

Gene conversion between *MT* alleles in shared genes is expected to reduce allelic differentiation. To test this prediction, we calculated *F*_*ST*_ over shared regions in 1 kb windows and examined its relationship with *Z*_*nS*_. We find a strong relationship (Fig. 3, *R*^2^ = 0.534*, p <* 2.2 *×* 10^*−*16^), suggesting recombination via gene conversion is an effective mode to homogenize genetic variation between *MT* alleles.

**Figure 3:**
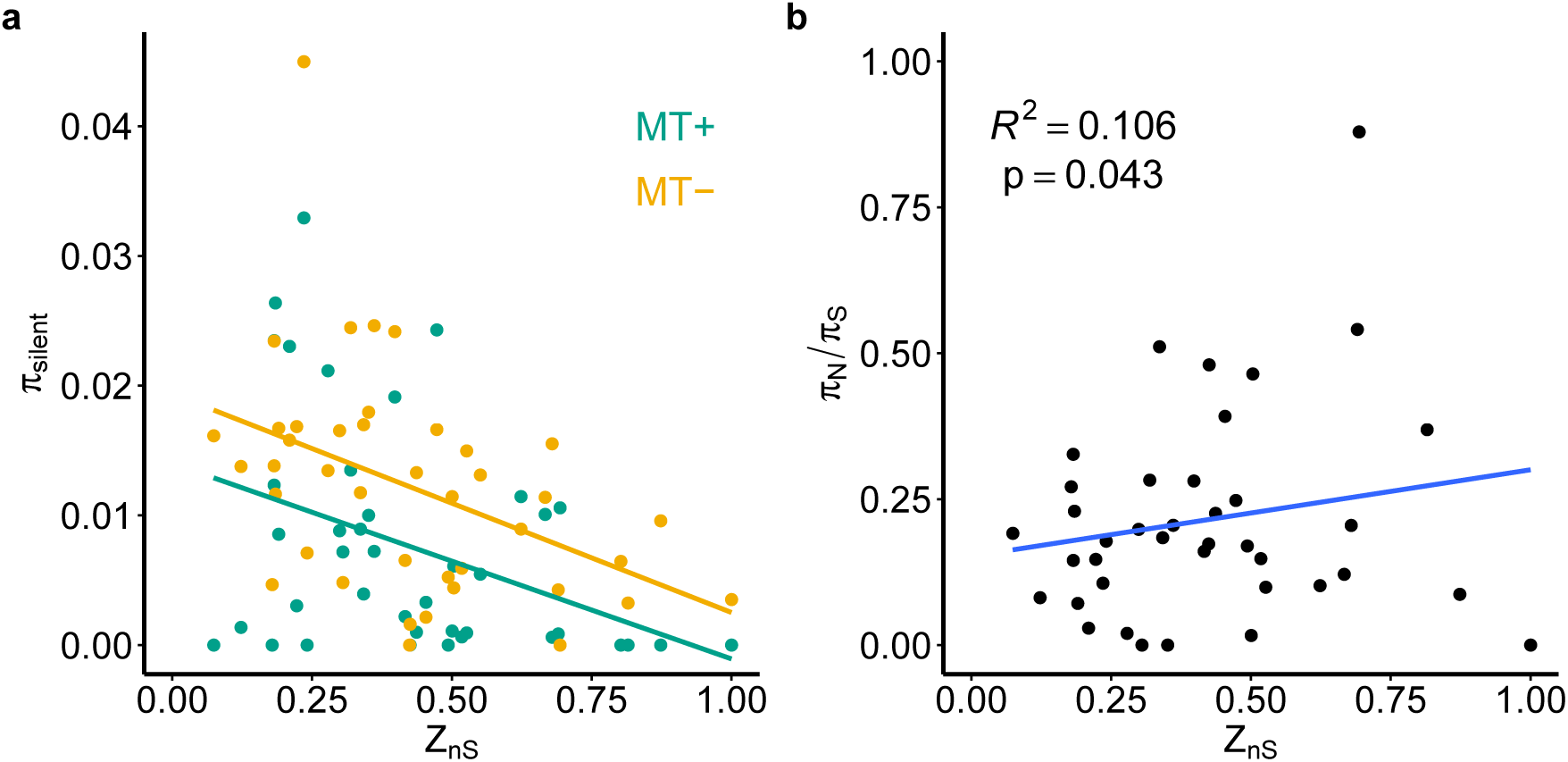
**a**. LD (*Z*_*nS*_) and neutral diversity at shared genes, grouped by mating type. **b**. *π*_*N*_ /*π*_*S*_ shows a weakly positive relationship with *Z*_*nS*_. In both plots, each point represents a gene with copies in both *MT* alleles.

### Genes with higher recombination rates exhibit higher selection efficacy and GC content

Recombination reduces interference between selected sites (Hill-Robertson effects) and thus improves the efficacy of selection to fix beneficial and purge deleterious mutations (Hill and Robertson, 1966, Felsenstein, 1974). Hill-Robertson effects result in a net reduction in linked neutral diversity due to either background selection or selective sweeps. Thus, we expect nucleotide diversity (*π*) in shared genes to be higher than that of *MT*-limited genes, which we found to be the case (*π* in shared genes = 0.0168, *pi* in *MT+* genes = 0.0037, *π* in *MT–* genes = 0.0022). We also find that *π* in each mating type correlates negatively with *Z*_*nS*_, suggesting that genes that undergo more recombination exhibit reduced effects of selection at linked sites (Fig. 3a; *MT+*: *R*^2^ = 0.134,*p* = 0.016; *MT–*: *R*^2^ = 0.187,*p* = 0.0047). Mean *π* in *MT* (*π* = 0.0168) is about half of autosomal diversity (*∼* 0.03, Flowers et al. 2015) though *MT+* diversity is about half of *MT–* diversity (*MT+ π* = 0.0065, *MT– π* = 0.0112). To then test whether genes with higher recombination rates undergo more efficient purifying selection, we calculated both the ratio of nonsynonymous to synonymous diversity (*π*_*N*_ /*π*_*S*_) as well as *Z*_*nS*_ in the coding regions of both shared and *MT*-limited genes in the *MT* locus. *π*_*N*_ /*π*_*S*_ almost always ranges from 0 to 1, with lower values suggesting more efficient selection; values above 1 are rare, and instead suggest that balancing selection may be acting to preserve nonsynonymous diversity.

Here, weighting by the number of sites in each gene, we find a weakly positive relationship between *Z*_*nS*_ and *π*_*N*_ /*π*_*S*_, suggesting that regions of higher recombination also show more efficient selection (Fig. 3b, *R*^2^ = 0.106, *p* = 0.043). This result excludes two genes, *MT0796* and *522915*, which showed extreme values of *π*_*N*_ /*π*_*S*_ (1.85 and 3.35 respectively); including these outlier genes makes the fit marginally nonsignificant (*R*^2^ = 0.094, *p* = 0.051) but the direction of the relationship remains the same.

We next examined whether recombination rate affected GC content in shared genes. Either or both of GC-biased gene conversion or selection on base composition are expected to drive increases in GC content in regions of higher recombination. Here, we calculated GC content at intronic, intergenic, and four fold degenerate sites (GC4) in 1 kb windows across shared regions, and found a significant but very weakly negative relationship between GC content and *Z*_*nS*_ (*R*^2^ = 0.023, *p* = 0.00028, slope = *−*0.048).

### Contrasting selection efficacy and GC-content evolution in *MT* with autosomal genes

To examine how selection efficacy in *MT*-limited and shared genes compares to that of autosomal genes, we then calculated *π*_*N*_ /*π*_*S*_ for 100 randomly sampled genes from chromosome 6 in addition to a further 100 randomly selected autosomal genes from across the remainder of the genome. Across the four annotations (*MT*-limited, shared, chromosome 6, and autosomal), a Kruskal-Wallis test showed no significant differences in *π*_*N*_ /*π*_*S*_ (*χ*^2^ = 2.623, *p* = 0.454) indicating that selection efficacy in shared *MT* genes is similar to that of non-*MT* genes. While this result also surprisingly suggests no significant difference in selection efficacy between *MT*-limited genes and autosomal genes, this comparison is likely underpowered by the fact that there are only 7 *MT*-limited genes in *MT* as compared to 51 shared *MT* genes included in our analysis.

We repeated the above analysis for GC4 across the four annotations of interest, but using our estimates of GC4 in 1 kb windows instead of genic values. Here, a Kruskal-Wallis test showed significant differences in GC4 (*χ*^2^ = 15.75, *p* = 0.0013). Subsequent paired Wilcoxon rank-sum tests showed that only differences in GC4 between shared genes and autosomal genes (both chromosome 6 and the randomly sampled dataset) were significantly different (*p* = 0.016 and *p* = 0.00098 for shared-chromosome 6 and shared-autosomal, respectively), with GC4 in shared genes higher in both cases. Interestingly, GC content levels for the *MT*-limited genes *MID* and *FUS1* were drastically lower than the remainder of the *MT* locus and the *C. reinhardtii* genome as a whole (*MID* GC4 = 0.526, *FUS1* GC4 = 0.396), despite the overall similarities seen in GC4 between shared and *MT*-limited regions (Table 1).

**Table 1:**
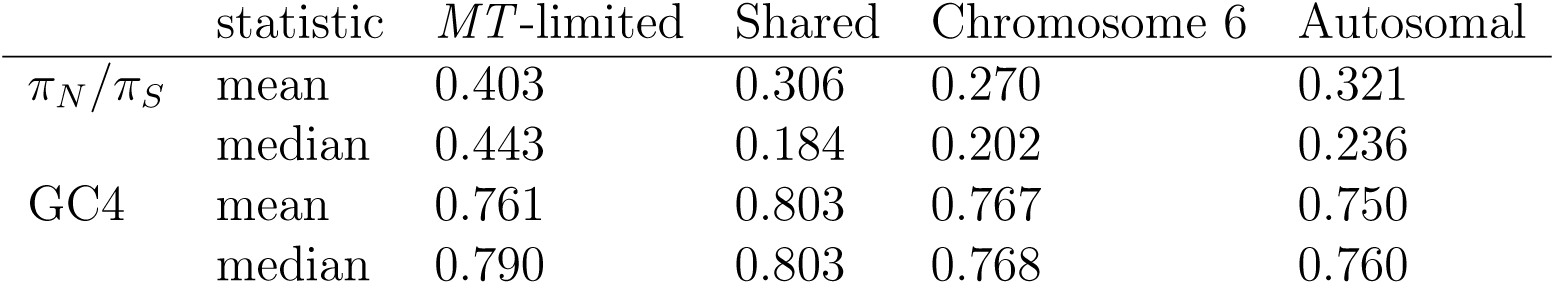
Summary of *π_N_ /π_S_* and GC content across annotations examined.

|  | statistic | *MT*-limited | Shared | Chromosome 6 | Autosomal |
| --- | --- | --- | --- | --- | --- |
| $\pi_N/\pi_S$ | mean | 0.403 | 0.306 | 0.270 | 0.321 |
|  | median | 0.443 | 0.184 | 0.202 | 0.236 |
| GC4 | mean | 0.761 | 0.803 | 0.767 | 0.750 |
|  | median | 0.790 | 0.803 | 0.768 | 0.760 |

### Breakdown of inter-chromosomal LD suggests biparental chloroplast inheritance

To test for evidence of leakage in uniparental organelle inheritance, we calculated pairwise LD (D’) between between SNPs from 1) the chloroplast genome and the *MT+* specific genes *fus* and *mta1*, 2) the mitochondrial genome and the *MT–* specific gene *mtd1*, 3) the *MT–* mitochondrial genome with itself, and 4) randomly sampled SNPs across all 17 chromosomes of *C. reinhardtii*. The distribution of obtained D’ values for these comparisons is shown in Figure 4.

**Figure 4:**
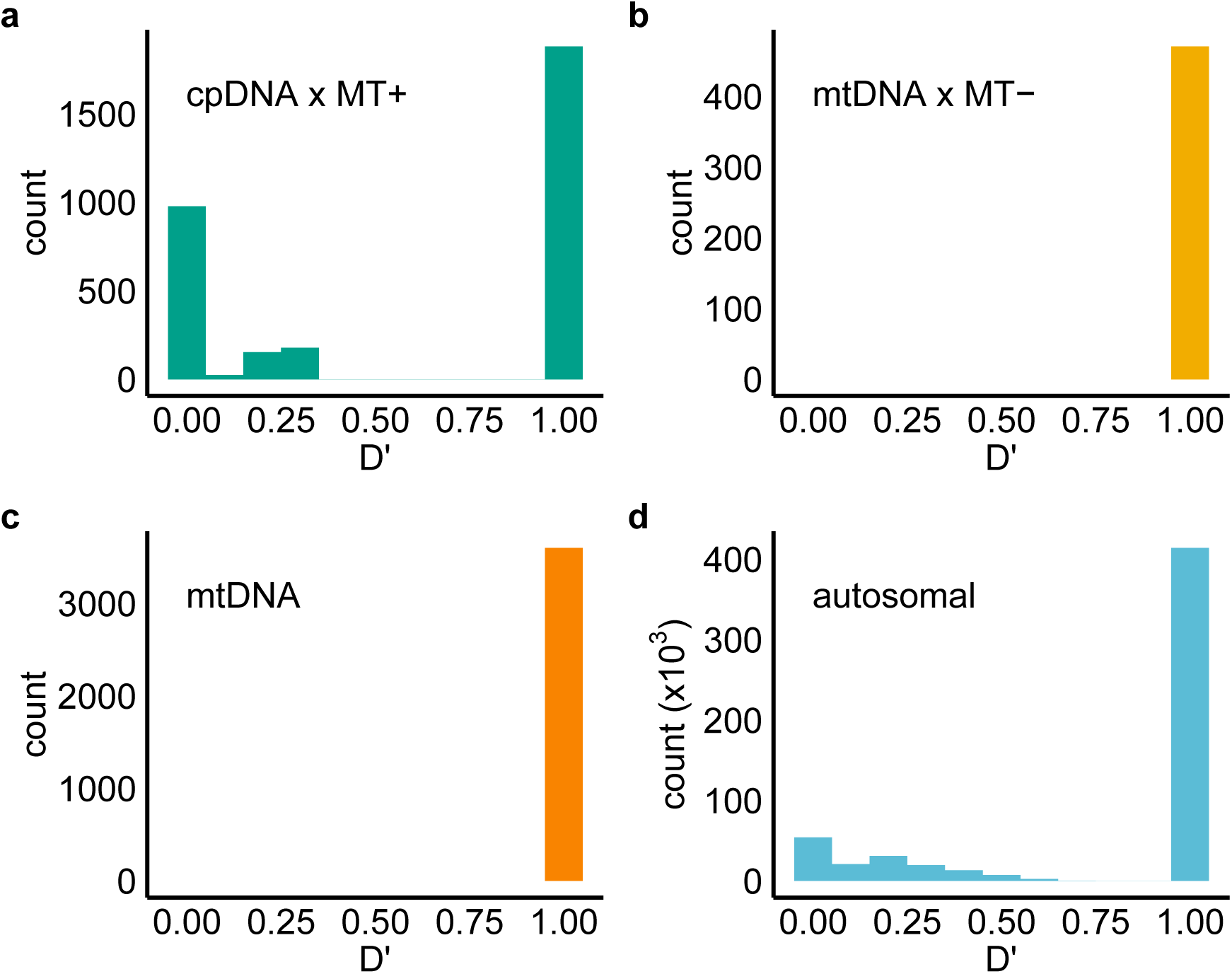
Distribution of D’ for SNPs across: **a**. cpDNA and *MT+*. **b** mtDNA and *MT–*. **c** The mitochondrial genome. **d**. Randomly drawn pairs of SNPs from across the genome.

LD between the chloroplast genome and corresponding *MT+* markers displays a large proportion of sites in complete linkage but also a clear pattern of LD breakdown (Fig. 4a; mean D’ = 0.611) thus implicating recombination between chloroplast genomes or some bi-parental inheritance. Interestingly, this appears to be an even lower level of inter-chromosomal LD than is observed in randomly selected pairs (Fig. 4d; mean D’ = 0.781). In contrast, the mitochondrial genome is in complete LD with the *MT–* limited genes (mean D’ = 1.0).

## Discussion

While many models of sex chromosome and mating type locus evolution require an initial supression of recombination, the evolutionary trajectory of mating type and sex determining regions may differ substantially. Previous work comparing the *MT* locus of *C. reinhardtii* with that of its anisogamous, multicellular relative *Volvox carteri* showed that *C. reinhardtii MT* alleles were two orders of magnitude less diverged by comparison, despite the two species sharing a homologous *MT* locus (Ferris et al., 2010). The subsequent finding that infrequent gene conversion was occurring between the two *C. reinhardtii MT* alleles provided an explanation for why the shared regions were not differentiated (De Hoff et al., 2013). Here, we examine patterns of coding and non-coding genetic diversity across the entire *MT* locus to estimate the extent of recombination rate variation and its consequences for selection on *MT* genes. We show that recombination rate predicts the degree to which shared regions have differentiated across *MT* haplotypes, and that genes with higher recombination rates exhibit greater selection efficacy against deleterious alleles. Furthermore, we use patterns of LD breakdown to show evidence for biparental inheritance of the chloroplast genome but not the mitochondrial genome.

Over the *MT* locus, we observe strongly elevated LD in the R domain, while LD in the flanking T and C domains resemble autosomal levels (Fig. 1). The pattern we show is expected, given that the T and C domains are highly syntenic while the R domain contains inversions and autosomal translocations (Ferris and Goodenough, 1994, Ferris et al., 2002, De Hoff et al., 2013), resulting in recombination suppression (Charlesworth, 1994, Wright et al., 2016). Although LD is elevated in the R domain, it is noteworthy that *Z*_*nS*_ is below 1 and comparable to other regions of chromosome 6 with high LD. It is in fact the absence of low LD windows in the R domain that differentiates it from the rest of the chromosome. Our hidden Markov model of recombination rate did not delineate a clear boundary at which recombination suppression begins on either bound of the R domain, similar to findings in the pseudo-autosomal regions of human sex chromosomes (Cotter et al., 2016). However, LD patterns in the T and C domains are indistiguishable from those of the surrounding autosomal regions. Although early evidence from laboratory crosses suggested the entirety of *MT* was recombination suppressed (Ferris and Goodenough, 1994), our results mirror those of De Hoff et al. (2013) in suggesting that both crossing over and non-crossover gene conversions may be occuring in these regions. Patterns of LD between *MT+* genes and the chloroplast genome also clearly support the existence of recombination and/or biparental inheritance in the chloroplast, in line with previous results (Boynton et al., 1987, Nakamura, 2010, Ness et al., 2016). In contrast, no such evidence exists for the *MT–* genes and the mtDNA, supporting strict uniparental inheritance of the mitochondria even over long evolutionary timescales.

Our results demonstrate that gene conversion has important consequences for the molecular evolution of *MT*. Firstly, we show that across the *MT* locus, shared regions with higher rates of recombination exhibit less differentiation between mating types (*F_ST_*, Fig. 2a), as would be expected in a program where gene conversion is widespread in the shared regions of the *MT* locus. Increased *F_ST_* in shared regions could result from both lower rates of gene conversion or selection to retain alleles with *MT*-specific selective effects. Although less selection on secondary sex characteristics is expected in isogamous systems, differential expression of shared genes between mating type alleles can still create selective pressure for allelic differentiation (Immler and Otto, 2015), and has been previously observed in the *C. reinhardtii MADS2* (De Hoff et al., 2013). Thus, the possibility that differential expression may be playing a role alongside gene conversion in determining differentiation levels cannot be excluded until allele-specific expression data across more shared genes are obtained. Finally, gene conversion across the *MT* locus is also expected to affect local GC content. Recent work on the strength of gBGC in *C. reinhardtii* found a significant GC bias in noncrossover gene conversions (Liu et al., 2017). Consistent with predictions from gBGC, we found that GC content was highest in shared *MT* genes, and that the *MT*-limited genes *MID* and *FUS1* had extremely low GC content.

We predicted that selection against deleterious mutations across the *MT* locus would be weaker in regions with less recombination due to interference between selected sites (Hill and Robertson, 1966, Felsenstein, 1974, Comeron et al., 2008). Indeed, we found that diversity levels were lower in shared genes with low recombination (Fig. 2a), while diversity in *MT*-limited genes was an order of magnitude lower than in shared genes. Both of these findings are consistent with background selection reducing neutral diversity. We see lower diversity in *MT+* than *MT–*, which may be reflective of previous reports that gene conversion in *MT* is biased in favour of *MT+* to *MT–* conversions (De Hoff et al., 2013), although the negative correlation between *π* and *Z*_*nS*_ is still present in both alleles (Fig. 3a). Furthermore, *MT* regions with higher LD exhibited higher *π*_*N*_ /*π*_*S*_ ratios, indicative of reduced purifying selection (Fig. 3b). Interestingly, the overall level of *π*_*N*_ /*π*_*S*_ in shared genes was not significantly different from those of autosomal genes, suggesting that gene conversion in shared genes may be enough to maintain equivalent levels of selection efficacy. With Hill-Robertson effects being a key mechanism in the degeneration of sex chromosomes (Bachtrog, 2013), this suggests that gene conversion is sufficient to facilitate effective purifying selection and prevent the accumulation of deleterious variants in *MT* in the absence of crossovers. Additionally, unlike crossovers, gene conversions can occur between inversions (Korunes and Noor, 2017), as previously observed in the R domain of *MT* (De Hoff et al., 2013). Similar patterns of periodic inter-chromosomal gene conversion for putatively adaptive purposes have been observed in the sex chromosomes of European tree frogs (Stöck et al., 2011), avian sex chromosomes (Wright et al., 2014), fungal mating type loci (Menkis et al., 2010, Sun et al., 2012), and in certain mammalian sex-linked orthologs (Pecon Slattery et al., 2000, Rosser et al., 2009, Peneder et al., 2017, Trombetta et al., 2017). However, the effect of gene conversion on selection efficacy in these regions has not yet been investigated.

Theoretical models predict reduced recombination between haploid sex chromosomes if different sexes undergo differing selective pressures (Immler and Otto, 2015), such as in anisogamous systems. The *MT* locus of *V. carteri*, for instance, is five times larger than *C. reinhardtii MT*, more differentiated, and also carries more sex-specific genes (Ferris et al., 2010). Differentiation between the *V. carteri MT* alleles likely occurred after the transition to anisogamy (Hiraide et al., 2013, Hamaji et al., 2018), suggesting that selection for recombination suppression followed the appearance of selective pressure on sex-specific traits. Thus, given our findings that gene conversion is a strong predictor of both selection efficacy and differentiation levels across *C. reinhardtii MT*, we argue reduced gene conversion plays an important role in the accumulation of heteromorphism and eventual degeneration in anisogamous systems. Taken together, our results suggest that in isogamous systems lacking secondary sexual characteristics, recombination plays an important role in reducing *MT* differentiation as well as degeneration through less efficient selection. Although suppressed recombination is necessary in many mating type loci to facilitate sexual reproduction and sequester sex-specific genes, there is less pressure to differentiate these regions than in the sex chromosomes of anisogamous systems, thus circumventing the degeneration characteristic of sex chromosomes. Evidence of degeneration is restricted to *MT*-limited genes in *C. reinhardtii*, which we show exhibit low diversity and less effective selection. In addition to *C. reinhardtii*, other isogamous algae also show very low levels of differentiation in shared regions (Hamaji et al., 2016, 2018, Coelho et al., 2018) suggesting that similar mechanisms are acting in the mating type loci of a broad array of species.

## Acknowledgements

Short read data are available at the European Nucleotide Archive under study accession ERP109393. This work was supported by a Natural Sciences and Engineering Research Council (NSERC) Discovery grant (RGPIN/06331-2016) and Canadian Foundation for Innovation John R. Evans Leaders fund (35591) to RWN. We thank S.I. Wright, A.M. Moses, and the Plant Evolutionary Genomics group at the University of Toronto for helpful discussions and suggestions. We also thank Brian Novogradac for facilitating computational support and HPCNODE1.

## Supplementary Information

**Table S1:** Field strains of *C. reinhardtii* used in this study. All strains were obtained from the *Chlamydomonas* Resource Center (`chlamycollection.org`). Mating types of *MT–* strains CC-3059 and CC-3062 are mislabelled as *MT+* on the Resource Center website, and are instead *MT–* individuals (R.J. Craig, personal communication)

| Strain | Collection Location/Year | Mating type |
| --- | --- | --- |
| CC-2936 | Farnham, Quebec/1993 | + |
| CC-2937 | Farnham, Quebec/1993 | + |
| CC-3060 | Farnham, Quebec/1993 | + |
| CC-3064 | Farnham, Quebec/1993 | + |
| CC-3065 | Farnham, Quebec/1993 | + |
| CC-3068 | Farnham, Quebec/1993 | + |
| CC-3071 | Farnham, Quebec/1993 | + |
| CC-3076 | MacDonald College, Quebec/1994 | + |
| CC-3086 | MacDonald College, Quebec/1994 | + |
| CC-2935 | Farnham, Quebec/1993 | - |
| CC-2938 | Farnham, Quebec/1993 | - |
| CC-3059 | Farnham, Quebec/1993 | - |
| CC-3061 | Farnham, Quebec/1993 | - |
| CC-3062 | Farnham, Quebec/1993 | - |
| CC-3063 | Farnham, Quebec/1993 | - |
| CC-3073 | Farnham, Quebec/1993 | - |
| CC-3075 | MacDonald College, Quebec/1994 | - |
| CC-3079 | MacDonald College, Quebec/1994 | - |
| CC-3084 | MacDonald College, Quebec/1994 | - |

